# CLEC5A and TLR2 are critical in SARS-CoV-2-induced NET formation and lung inflammation

**DOI:** 10.1101/2022.02.01.478701

**Authors:** Pei-Shan Sung, Shao-Ping Yang, Yu-Chun Peng, Cheng-Pu Sun, Mi-Hwa Tao, Shie-Liang Hsieh

**Author notes:** **Corresponding author:** Shie-Liang Hsieh, Distinguished Research Fellow, Genomics Research Center, Academia Sinica, 128 Academia Road, Sec. 2, Nankang District, Taipei 115, Taiwan.

## Abstract

Coronavirus-induced disease 19 (COVID-19) infects more than three hundred and sixty million patients worldwide, and people with severe symptoms frequently die of acute respiratory distress syndrome (ARDS). Autopsy demonstrates the presence of thrombosis and microangiopathy in the small vessels and capillaries. Recent studies indicated that excessive neutrophil extracellular traps (NETs) contributed to immunothrombosis, thereby leading to extensive intravascular coagulopathy and multiple organ dysfunction. Thus, understanding the mechanism of severe acute respiratory syndrome coronavirus 2 (SARS-CoV-2)-induced NET formation would be helpful to reduce thrombosis and prevent ARDS. It has been shown that sera from individuals with COVID-19 triggered NET release in vitro, and spleen tyrosine kinase (Syk) inhibitor R406 inhibited NETosis caused by COVID-19 plasma. However, the serum components responsible for NET formation are still unknown. In this study, we found that virus-free extracellular vesicles (EVs) from COVID-19 patients (COVID-19 EVs) induced robust NET formation via Syk-coupled C-type lectin member 5A (CLEC5A). Blockade of CLEC5A inhibited COVID-19 EVs-induced NETosis, and simultaneous blockade of CLEC5A and TLR2 further suppressed SARS-CoV-2-induced NETosis in vitro. Moreover, thromboinflammation and lung fibrosis were attenuated dramatically in *clec5a^-/-^/tlr2^-/-^* mice. These results suggest that COVID-19 EVs play critical roles in SARS-CoV-2-induced immunothrombosis, and blockade of CLEC5A and TLR2 is a promising strategy to inhibit SARS-CoV-2-induced intravascular coagulopathy and reduce the risk of ARDS in COVID-19 patients.

## Introduction

SARS-CoV-2 is the etiological agent of the COVID-19 [1, 2], and has claimed more than 5.62 million lives worldwide. One of the clinical features of severe SARS-CoV-2 infection is extensive infiltration of neutrophils and macrophages in lung, and COVID-19 patients suffer from ARDS caused by extensive lung inflammation, tissue injury, and fibrosis [3]. In addition, pulmonary embolism [4, 5], microangiopathy, cerebral infarction, [6] and visceral organ thrombo-emboli are frequently observed in post-mortem autopsy of COVID-19 patients [7, 8].

Even though the pathogenesis of thrombo-emboli formation in COVID-19 patients is still unclear, recent studies indicate that excessive NET production is associated with thrombo-emboli formation in human diseases. It has been reported that elevated levels of cell-free DNA, myeloperoxidase, and citrullinated histone H3 are noted in the sera of COVID-19 patients, and higher levels of NET formation correlated with disease severity [9, 10]. Thus, targeting excessive NET formation is speculated to reduce pulmonary inflammation and thrombosis in COVID-19 patients [11].

Because platelets are hyperactivated in critically ill COVID-19 patients [12, 13], and platelets-neutrophil interactions play critical roles in endothelial damage and immunothrombosis of COVID-19 patients [14–16], we are interested to understand the molecular mechanism of platelets-mediated enhancement of NET formation and pulmonary inflammation in SARS-CoV-2 infection. Recently, SARS-CoV-2 was shown to activate platelets to enhance thrombosis in vivo [17], and serum levels of platelet-derived EVs (PLT-EVs) strongly associated with severity of SARS-CoV-2 infection [18]. Moreover, sera from COVID-19 patients trigger NET formation in neutrophils isolated from healthy donors [19]. These observations suggest that SARS-CoV-2 may activate platelets to release EVs, thereby induce NET formation and immunothrombosis in COVID-19 patients.

We have demonstrated that dengue virus (DV) [20–23] and microbes [24] bind and activate CLEC5A and TLR2 to release proinflammatory cytokines and promote NET formation. Moreover, DV activates Syk-coupled C-type lectin member 2 (CLEC2) in platelets to release EVs to enhance DV-induced NET formation and macrophage activation via CLEC5A and TLR2 [25, 26]. Recently, SARS-CoV-2 spike protein was reported to induce NETosis in Syk-dependent manner [27], suggesting that C-type lectins might play important roles in SARS-CoV-2-induced NET formation. Moreover, TLR2 was shown to sense SARS-CoV-2 membrane protein to activate NF-kB in monocyte and macrophages [28, 29]. Because CLEC5A and TLR2 are associated and co-activated by platelet-derived EVs (PLT-EVs) [25], we asked whether CLEC5A and TLR2 also contributed to SARS-CoV-2-induced immunothrombosis.

Here we report that COVID-19 EVs express abundant markers of activated platelets, and induce NET formation via CLEC5A and TLR2. Simultaneous blockade of CLEC5A and TLR2 inhibited SARS-CoV-2-induced NET formation in vitro, and thromboinflammation and fibrosis were dramatically attenuated in *clec5c^-/-^/tlr2^-/-^* mice. These observations suggest that EVs from virus-activated platelets play as endogenous danger signals to trigger NETs and inflammatory reactions via CLEC5A and TLR2, and blockade of CLEC5A and TLR2 may become a promising strategy to attenuate SARS-CoV-2-induced thrombus and coagulopathy in COVID-19 patients in the future.

## Materials and methods

### Reagents and antibodies

All culture media and buffers were purchased from Gibco. The source of anti-hCLEC5A mAb (clone 3E12A2) and anti-hTLR2 mAb (# MAB2616, R and D system) were as previously described^18^. Antibodies for immunofluorescence staining are as followings: rabbit anti-citrullinated histone H3 (#NB100-57135; Novus), goat anti-human/mouse myeloperoxidase polyclonal antibody (# AF3667, R&D system). Antibodies for immunohistochemical (IHC) staining are: rabbit anti-citrullinated histone H3 (#NB100-57135; Novus), goat anti-human/mouse myeloperoxidase polyclonal antibody (#AF3667, R&D system), anti-CD11b antibody (#ab13357, Abcam), anti-CD64 antibody (#MA5-29704, Invitrogen), anti-Siglec-F antibody (#PA5-11675, Invitrogen), anti-F4/80 antibody (#ab74383, Abcam), anti-CCR2 antibody (#NBP-35334, Novus Biologicals), anti-Ly6C antibody (#SC-23080, Santa Cruz). Secondary antibodies: donkey anti-mouse IgG (H+L) Alexa 488-conjugated antibody (#715-545-151, Jackson ImmunoResearch), donkey anti-goat IgG (H+L) Alexa 647-conjugated antibody (#705-605-147, Jackson ImmunoResearch), donkey anti-Human IgG (H+L) HRP-conjugated antibody (#709-035-149, Jackson ImmunoResearch), HRP-conjugated donkey anti-rabbit IgG (H+L) (#711-035-152, Jackson ImmunoResearch), HRP-conjugated donkey anti-goat IgG (H+L) (#SC-2020, Santa Cruz), HRP-conjugated donkey anti-mouse IgG (H+L) (#715-035-150, Jackson ImmunoResearch).

### Isolation of human primary neutrophils and platelets

Blood was drawn from healthy donors (unvaccinated with SARS-CoV-2 vaccine) into an anticoagulant ACD-containing syringe (ACD: blood ratio = 1:6, v/v), platelet-rich plasma was collected by centrifuge at 230 × g for 15 min. Pellet of platelets was harvested by centrifugation at 1000 × g for 10 min, then suspended in Tyrode’s buffer. For human neutrophils, whole blood was laid on the Ficoll-Paque (GE Healthcare, 45-001-748) and centrifuged at 500 × g for 15 min to get red blood cells (RBCs)-granulocytes-rich layer. After RBC lysis, neutrophils were washed and suspended in RPMI containing 10 % serum from unvaccinated blood type A/B healthy donors. The protocol was approved by the Human Subject Research

Ethics, Academia Sinica (AS-IRB-BM-20025).

### Isolation of mouse primary neutrophils and platelets

Platelets-rich plasma from murine peripheral blood was collected into ACD-containing Eppendorf and centrifugated at 230 × g for 4 min. Platelets were washed once and harvested by centrifugation at 1000 × g for 4 min, pellet was suspended in Tyrode’s buffer. For neutrophil isolation, bone marrow was collected and incubated with RBC lysis buffer. Bone marrow cells were suspended in a 45 % Percoll solution, then laid on 52 %, 63 %, and 81 % Percoll, and were further centrifugated at 1000 × g for 30 min. Neutrophils were harvested from layer 3 of the Percoll gradient, followed by washing with HBSS twice before being suspended in RPMI containing10 % FBS.

### SARS-CoV-2 propagation

SARS-CoV-2 Taiwan/4/2020 was propagated in Vero E6 cells. Viral titer was determined by observation of the cytopathic effect (CPE) in Vero cells. This strain was used in all of the experiments.

#### Production and purification of pseudotyped lentivirus

The pseudotyped lentivirus carrying the SARS-Co-2 spike protein was generated as described previously [30]. In brief, HEK-293T cells were transiently transfected with pLAS2w.Fluc.Ppuro, pcDNA3.1-2019-nCoV-S, and pCMV-Δ R8.91 using TransITR-LT1 transfection reagent (Mirus). Cell debris was removed by centrifugation at 4,000×g for 10 min, followed by passing the supernatant through a 0.45 μm syringe filter (Pall Corporation). For pseudotyped virus purification and concentration, supernatant was mixed with 0.2× volume of 50 % PEG 8,000 (Sigma) and incubated at 4 °C for 2 h. The pseudotyped lentivirus was then recovered by centrifugation at 5,000×g for 2 h, and resuspended in sterilized phosphate-buffered saline, aliquoted, and stored at −80 °C.

### Mouse model for SARS-CoV-2 infection

Virus preparation and inoculation of SARS-CoV-2 into C57BL/6 and *clec5a^-/-^tlr2^-/-^* mice were as described [31]. Lung tissue was collected at 3 days and 5 days post-infection for further analysis. All the animal experiments followed the protocol approved by the Institutional Animal Care and Use Committee (IACUC) at AS core (protocol ID 20-10-1521).

### Collection of tissues for RNA isolation

Mice were sacrificed at 3 days and 5 days post-infection. For RNA isolation, lung was dug into TRizol-containing MagNA Lyser Green Beads (Roche) for tissue homogenization and further isolated RNA using TriRNA Pure Kit (Geneaid) according to the manufacturer’s instruction. cDNA was synthesized using the RevertAid First Strand cDNA Synthesis Kit and the real-time PCR was performed as followed condition: 95 °C for 5min, followed by 30 cycles of 15 s at 95 °C, 30 s at 58 °C, and 30 s at 72 °C. The primer sequences were listed in Supplementary Table I. Data were shown as fold change compared to mock after normalized to GAPDH.

### Immunohistochemistry (IHC)

Lung tissue was fixed in 10 % paraformaldehyde for 48 h and embedded in paraffin subsequently. Tissue sections were deparaffined and rehydrated before multiple-color fluorescent staining using Opal 7-Color IHC Kits (Akoya bioscience). Samples were incubated with primary antibody (1:50) at 4°C overnight, followed by incubation with secondary antibody (1:100) at room temperature for 1 h. The Opal fluorescent dye was applied according to the vendor’s instructions. Images were captured with a Leica confocal microscope with white light laser system (TCS SP8X-FALCON) and exported using the Leica Application Suite X software. The NET/thrombosis quantitation and cell population analysis by MetaMorph^TM^ image software.

### Collagen deposition

Lung sections were de-paraffined and re-hydrated before being stained with Picro Sirius Red Stain Kit (#ab150681, Abcam), and images were captured by a light microscope with polarized light (Nikon). Quantification of collagen was performed by MetaMorph^TM^, and the level of collagendeposition was presented as area (μm^2^) of collagen under 40 × magnification.

### Isolation of extracellular vesicles (EVs)

Plasma from healthy donors and COVID-19 patients with severe pneumonia were centrifugated at 3500 × g for 15 min to remove cells and debris. Supernatants were further centrifuged at 100,000 × g for 1.5 h at 4 °C. Pellets were washed with saline and centrifuged at 100,000 × g for 1.5 h at 4 °C, followed by resuspension in 1 ml of saline. The protein concentration of EVs was determined by DC protein assay (Bio-Rad) according to the manufacturer’s instruction.

### Induction of neutrophils extracellular traps (NETs)

Human neutrophils (4×10^5^/ml) were seeded on poly-L-lysine-coated 12 mm coverslip in 24 well and simultaneously incubated with SARS-CoV-2 (MOI=0.1 or 1) in presence of autologous platelets (4 × 10^6^/ml) for 5 or 20 h at 37 °C. For SARS-CoV-2-spike pseudotyped virus stimulation, human neutrophils (4×10^5^/ml) were incubated with SARS-CoV-2-spike pseudotyped virus (MOI=0.25) and co-incubated with autologous platelets (4 × 10^6^/ml) for 3 h at 37 °C. For blocking assay, neutrophils were preincubated with isotype (100 μg/ml), anti-CLEC5A mAb (100 μg/ml, clone 3E12A2), anti-TLR2 mAb (100 μg/ml, R&D system), or a mixture of anti-CLEC5A mAb and anti-TLR2 mAb for 30 min at room temperature before incubation with SARS-CoV-2. For EVs stimulation assay, neutrophil was incubated with EVs from the plasma of healthy controls (HC-EVs) or COVID-19 EVs for 3 h at 37 °C.

### Visualization and quantification of NET structure

Cells were immersed in fixation buffer (containing 4 % paraformaldehyde) overnight, followed by permeabilization using 0.5 % Triton X100 in PBS, then incubated with anti-MPO antibody (1:100), anti-citrullinated histone antibody (1:100), and Hoechst 33342 (1:100000). The level of NETs was calculated using the histone image captured by a Leica confocal microscope with white light laser system (TCS SP8 X-FALCON), and analyzed by MetaMorph^TM^ software.

### Mass spectrometry analysis

Mass spectrometry analysis was performed in the Mass Spectrometry Core Facility located in the Genomic Research Center, Academia Sinica. In brief, EVs samples were lysed by RIPA solution containing phosphatase and protease inhibitors. Before the mass spectrometry analysis, samples were washed in PBS, followed by trypsin digestion before subjected to LTQ Orbitrap XL mass spectrometer (Thermo Fisher Scientific Inc.). Data were further analyzed by the Ingenuity Pathways Analysis (IPA) software.

## Results

### SARS-CoV-2 induces robust NET formation in the presence of platelets

To understand the role of platelet in SARS-CoV-2-induced NET formation, SARS-CoV-2 were incubated with human neutrophils in the presence or absence of autologous platelets. At 5 h post-incubation with SARS-CoV-2 (MOI=1) (2^nd^ panel from left, Fig. 1a), SARS-CoV-2 alone induced colocalization of citrullinated histone (Cit-H3), chromosomal DNA, and myeloperoxidase (MPO) within neutrophils. In contrast, robust aggregated NETs were observed in the presence of platelets (4^th^ panel from the left, Fig. 1a), which is distict from the thread-like NET structure induced by DV/platelets [25]. The detail of NET structure was presented as separated panels in Supplementary Figure. 1, and the level of NETs was measured by Cit-H3 area (mm^2^) (Fig. 1b). These observations demonstrated the critical role of platelets in SARS-CoV-2-induced NET formation.

**Figure 1.**
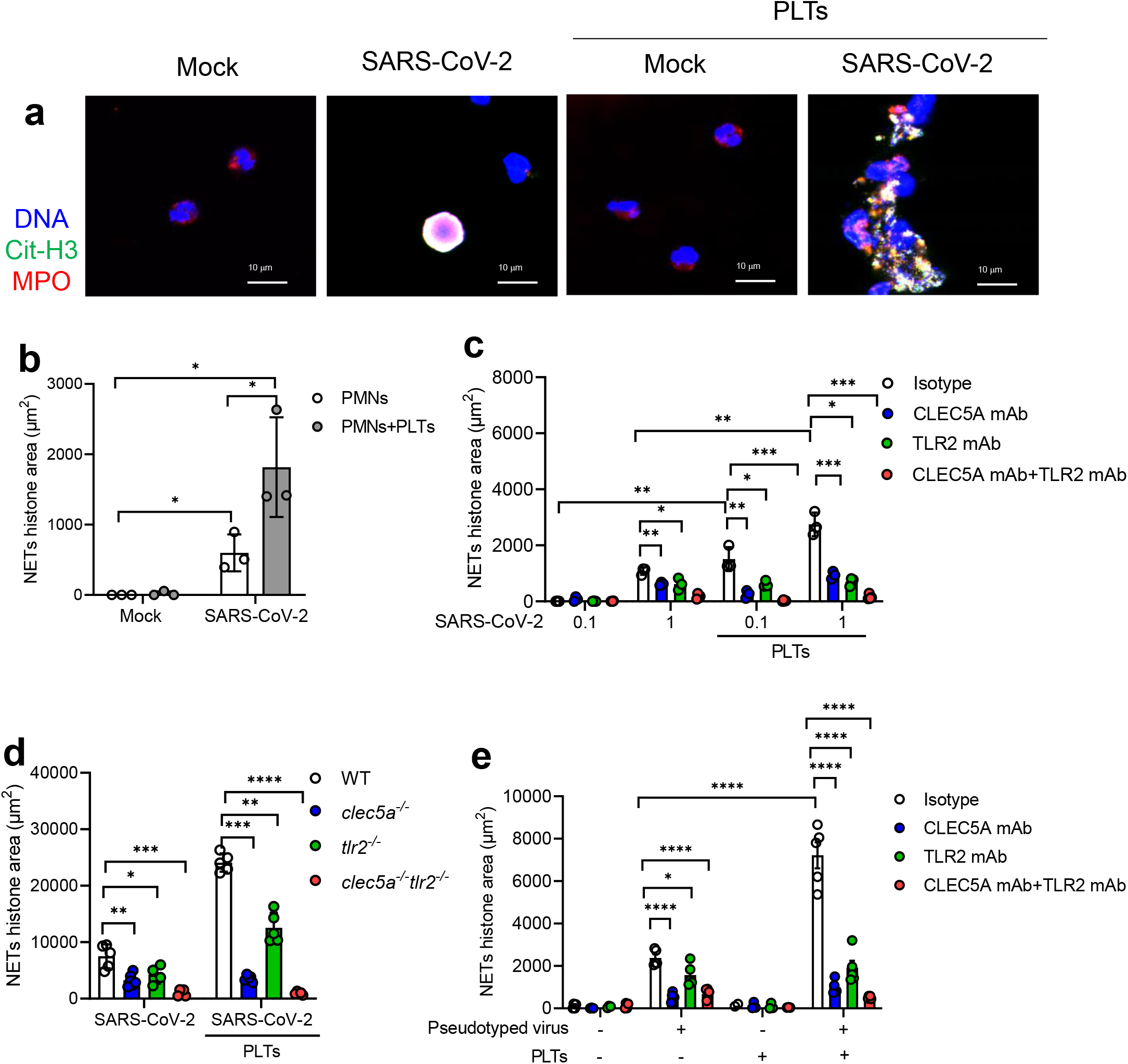
CLEC5A and TLR2 are responsible for SARS-CoV-2-induced NET formation. **(a&b)** Neutrophils (4 × 10^5^/ml) from healthy donors were incubated with SARS-CoV-2 (MOI = 1) with or without autologous platelets (4 × 10^6^/ml) for 5 h at 37 °C. The scale bar is 10 μm. The detailed structure of SARS-CoV-2-induced NET formation was observed under a confocal microscope (Leica). NET formation was visualized by fluorescent staining of DNA (blue), histone (green), and MPO (red) **(a)**. NETs level was measured by MetaMorph software and presented as Cit-H3 area (mm^2^) **(b)**. **(c)** Human neutrophils (4 × 10^5^/ml) were pretreated with anti-hCLEC5A mAb (3E12A2, 100 μg/ml), anti-TLR2 mAb (# MAB2616, 100 μg/ml), or combination of both antibodies for 30 min at room temperature, followed by incubation with SARS-CoV-2 (MOI = 0.1 and 1) in the presence or absence of platelets (4 × 10^6^/ml) for 5 h and 20 h. The level of NET formation was determined by histone area (μm^2^). **(d)** Neutrophils (4 × 10^5^/ml) from WT, *clec5a^-/-^tlr2^-/-^,* and *clec5a^-/-^tlr2^-/-^* mice were incubated with SARS-CoV-2 (MOI = 1) in the presence or absence of WT platelets (4 × 10^6^/ml) for 5 h at 37 °C. **(e)** Human neutrophils were pre-treated with anti-hCLEC5A mAb (3E12A2, 100 μg/ml), anti-TLR2 mAb (# MAB2616, 100 μg/ml), or combination of both antibodies for 30 min at room temperature, followed by incubation with SARS-CoV-2 spike pseudotyped virus (MOI = 0.1) in the presence or absence of autologous platelets (4 × 10^6^/ml) for 3 h. Data are mean ± SEM and repeats of 3 to 5 independent experiments. *p<0.05, **p < 0.01, ***p < 0.001, ****p < 0.0001 (Student’s t-test).

### CLEC5A and TLR2 are critical in NET formation caused by SARS-CoV-2

Our previous works demonstrated that CLEC5A and TLR2 were critical for virus and microbes-induced NET formation [25, 26], thus we further asked whether CLEC5A and TLR2 contributed to SARS-CoV-2-induced NETosis. We found that SARS-CoV-2 alone induced NET formation in a higher dose (MOI=1) at 5 h post-incubation (left, Fig. 1c). In the presence of platelets, SARS-CoV-2-induced NET formation was further enhanced (right, Fig. 1c). While NET formation was partially inhibited by anti-CLEC5A mAb and anti-TLR2 mAb, simultaneous blockade of CLEC5A and TLR2 almost completely suppressed SARS-CoV-2-induced NET formation (Fig. 1c). This observation suggests that platelet is a potent enhancer in SARS-CoV-2-induced NET formation.

We further incubated SARS-CoV-2 and SARS-CoV-2 plus platelets (SARS-CoV-2/platelets) with wild-type (WT) and *clec5a^-/-^tlr2^-/-^* neutrophils, respectivley, in the absence (left, Fig. 1d) or presence of WT platelets (right, Fig. 1d). We found that SARS-CoV-2 and SARS-CoV-2/platelets-induced NET formation was partially attenuated in *clec5a^-/-^* neutrophils and *tlr2^-/-^* neutrophils, while NET formation was almost undetectable in *clec5a^-/-^* neutrophils. This observation further confirms the critical roles of CLEC5A and TLR2 in SARS-CoV-2 and SARS-CoV-2/platelets-induced NET formation.

We further asked whether SARS-CoV-2 spike protein contributed to platelet activation by incubating SARS-CoV-2 spike pseudotyped virus with neutrophils for NETosis assay. We found that platelets still enhanced SARS-CoV-2 spike pseudotyped virus-induced NET formation, though the enhancing effect was less obvious than SARS-CoV-2. Blockade of TLR2 (green bar) and CLEC5A (blue bar) partially inhibited NET formation, simultaneous blockade of CLEC5A and TLR2 abolished SARS-CoV-2 spike pseudotyped virus-induced NET formation dramatically (Fig. 1e). This observation suggests that SARS-CoV-2 spike protein co-activates CLEC5A and TLR2 to induce NET formation.

### COVID-19 EVs induced NET formation via CLEC5A

As COVID-19 sera was reported to induce NET formation [19], and serum levels of platelet-derived EVs (PLT-EVs) correlated with disease severity [18], we asked whether COVID-19 EVs induced NET formation was via CLEC5A and TLR2. Firstly, we compared the protein components of EVs isolated from serum samples of COVID-19 patients (COVID-19 EVs) and normal individuals (HC-EVs) by mass spectrometry, and data were analyzed by ‘Ingenuity Pathway Analysis’ (IPA, QIAGEN) software. We found that molecules involved in platelet degranulation (CD9, platelet factor 4 (PF4)), aggregation, and activation [CD9, CLEC1B (also known as CLEC2)] were upregulated dramatically in COVID-19 EVs, while none of these proteins were detectable in EVs from healthy donors (Fig. 2a). In addition to these upregulated molecules, proteins specifically expressed in COVID-19 EVs were listed in Table I. We further confirmed the IPA results by flow cytometry analysis. We found that platelet activation markers (CD41a/b, CD62p), integrins (CD9, CD29, CD49e), adhesion molecules (CD31), and other activation marker (CD45, CD69) were upregulated (Fig. 2b & 2c). We then incubated neutrophils with HC-EVs and COVID-19 EVs, respectively, to compare their abilities to induce NET formation. While EVs from healthy control were unable to induce NET formation, COVID-19 EVs induced robust NET formation, which was blocked efficiently by anti-CLEC5A mAb, but not anti-TLR2 mAb (Fig. 2d). This observation suggests that COVID-19 EVs have potent activity to induce NET formation via CLEC5A.

**Figure 2.**
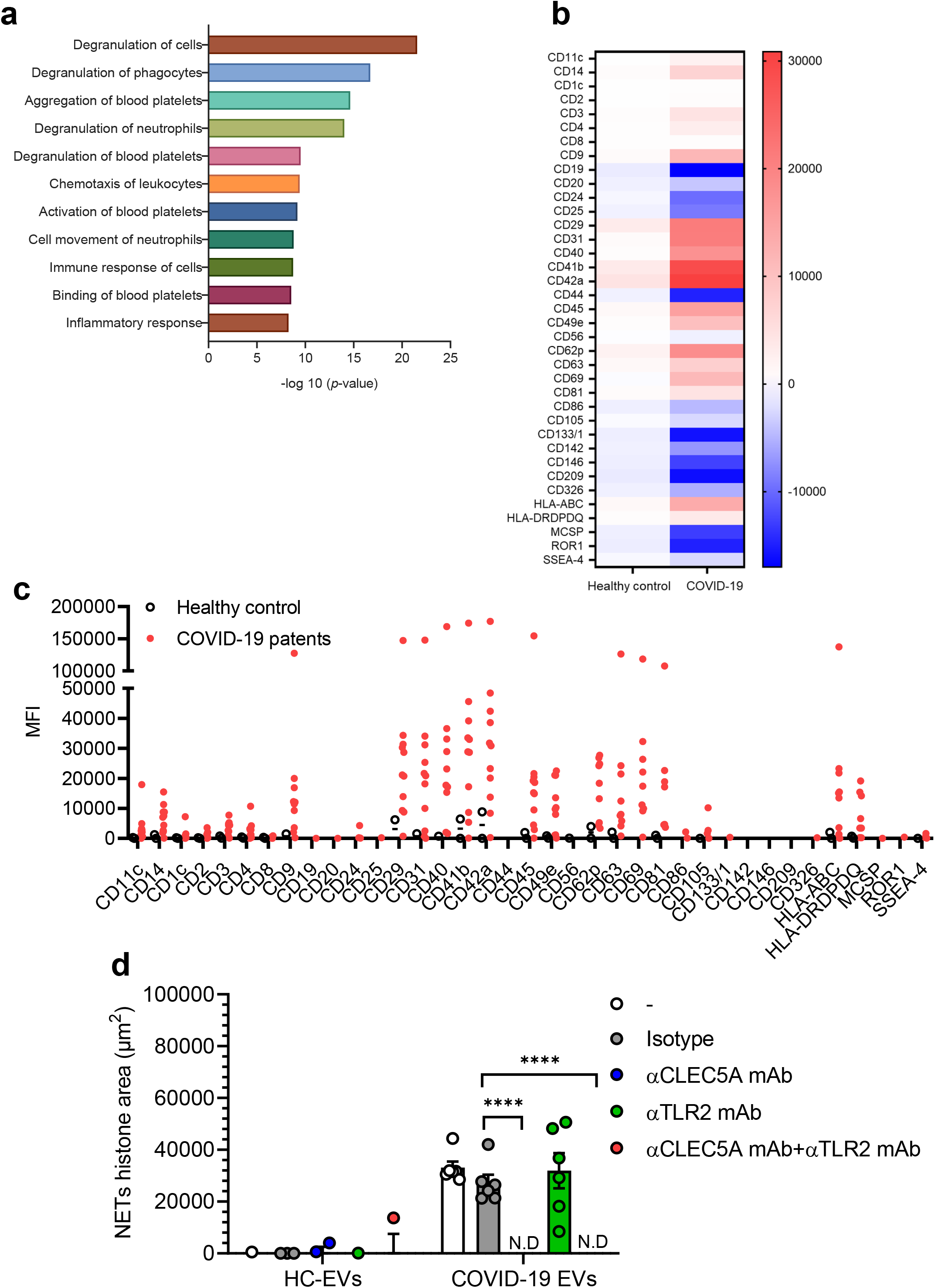
COVID-19 EVs are derived from activated platelets and induce NET formation via CLEC5A. **(a)** EVs from healthy controls (HCs-EVs, n=5) and COVID-19 patients (COVID19-EVs, n=5) were harvested by ultracentrifugation, then lysed in RIPA solution before subjected to mass spectrometry analysis. Proteins expressed in COVID-19 EVs, but not in HCs EVs, were further analyzed using the QIAGEN Ingenuity Pathway Analysis (QIAGEN IPA) software. Proteins which were expressed in all the COVID19-EVs were displayed. **(b&c)** HCs-EVs (n=10) and COVID19-EVs (n=10) were analyzed by flow cytometry, and markers highly activated in COVID-19 platelets were expressed as a heat map (b) or by mean fluorescence intensity (c)**. (d)** Neutrophils were pre-incubated with anti-CLEC5A mAb (3E12A2, 100 μg/ml), anti-TLR2 mAb (# MAB2616, 100 μg/ml), or both anti-CLEC5A mAb (3E12A2, 100 μg/ml) and anti-TLR2 mAb (# MAB2616, 100 μg/ml), for 30 min at room temperature, followed by incubation with EVs (1 μg/ml) from COVID-19 patients (n=6) at 37°C for 3 h. Data are mean ± sd and repeats of at least three independent experiments. \**p*<0.05, *\*\*p* < 0.01, *\*\*\*p* < 0.001, *\*\*\*\*p* < 0.0001 (Student’s *t*-test).

### CLEC5A and TLR2 are critical in SARS-CoV-2-induced thromboinflammation

We have used recombinant adeno-associated virus to introduce human ACE2 (hACE2) into wild type mice to establish a SARS-CoV-2 infection model system [31]. To evaluate the roles of CLEC5A and TLR2 in SARS-CoV-2-induced thromboinflammation, the AAV-hACE2 was introduced to wild-type littermates (WT) and *clec5a^-/-^tlr2^-/-^* mice to address this question. In WT mice, SARS-CoV-2 upregulated the expression of IL-6, IFN-γ, CCL-2, and IP-10 dramatically (more than 10 folds), while TNF-α, IL-1ß, IL-10, and chemokines (CXCL1, CXCL2, CXCL5) were also upregulated (Fig. 3a). Compared to WT mice, the expression of proinflammatory cytokines and chemokines were downregulated in *clec5a^-/-^tlr2^-/-^* mice. These observations demonstrated the critical role of CLEC5A and TLR2 in SARS-CoV-2-induced inflammatory reactions in lung tissues. We further detected NET formation in lung tissues after SARS-CoV-2 infection by examining the localization of DNA (blue), myeloperoxidase (green), citrullinated histone H3 (red), and platelet marker CD42b (yellow)(Fig. 3b). We found intense NET formation in lung tissue at day 3 and day 5 post-SARS-CoV-2 infection, while NET formation was almost undetectable in *clec5a^-/-^tlr2^-/-^* mice (Fig. 3b). The NET area and thrombi were further quantified by using anti-MPO antibody (Fig. 3c) and anti-CD42b antibody (Fig. 3d).

**Figure 3.**
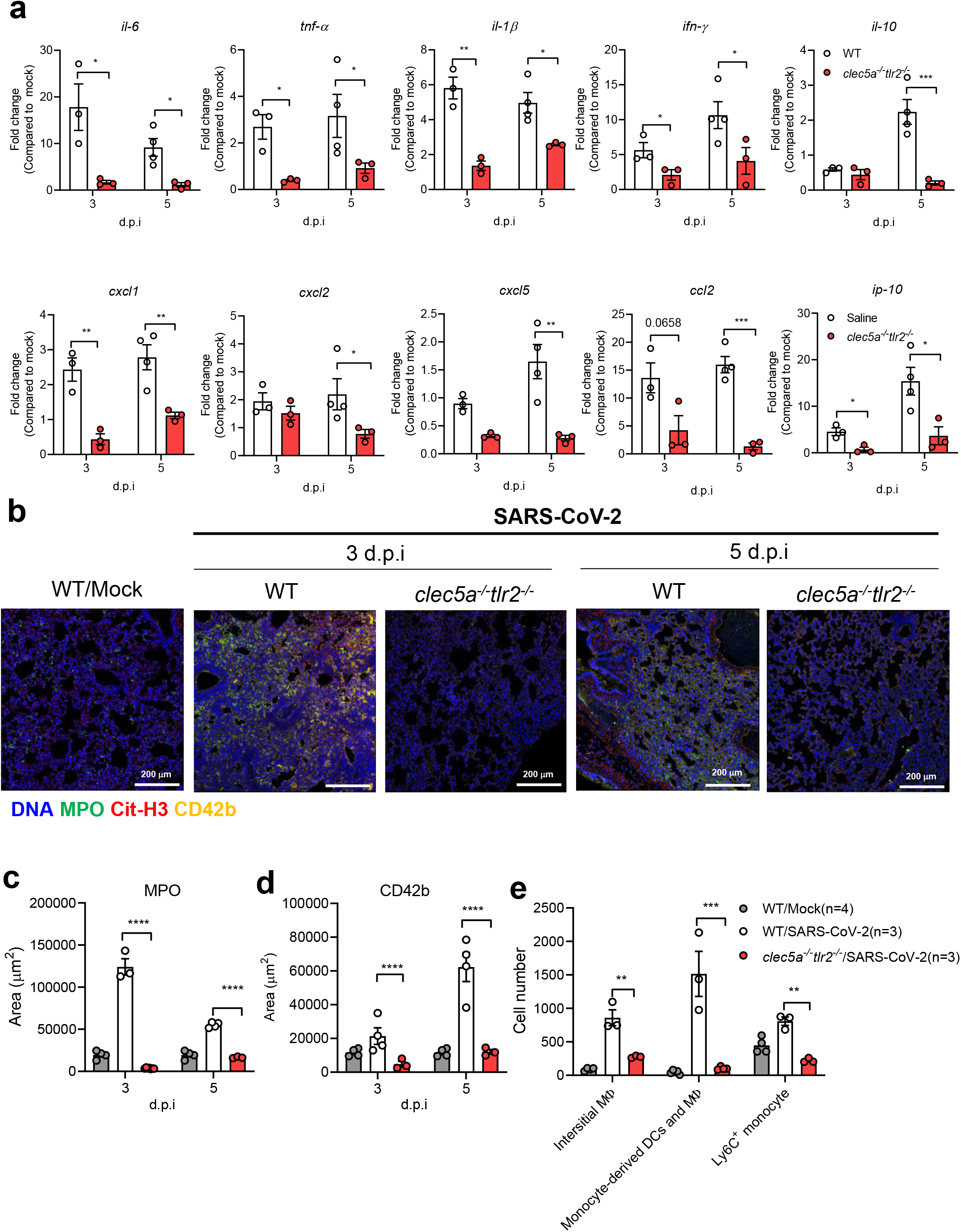
Attenuation of SARS-CoV-2-induced thromboinflammation in CLEC5A and TLR2 deficient mice. C57BL/6 mice (WT) (n=3) and *clec5a^-/-^tlr2^-/-^* mice (n=3)were inoculated with AAV-hACE2 for 14 days, followed by intranasal inoculation of SARS-CoV-2 (8 × 10^4^ PFU/per mice). Tissues were collected at 3 days and 5 days post-infection. **(a)** The level of proinflammatory cytokines and chemokines were measured by real-time PCR and presented as fold change (compared to AAV-hACE2 uninfected mice/mock). (**b-d)** NET structure and thrombus were detected by Hoechst. 33342 (blue), anti-MPO antibody (green), anti-citrullinated histone H3 (red), anti-CD42b antibody (yellow) **(b)**, and images were captured by a confocal microscope and subjected to determine the area of MPO **(c)** and CD42b **(d)** using MetaMorph^TM^ software. **(e)** Cell infiltrated to lung. Interstitial macrophage (interstitial MΦ) was defined as CD11b^+^CD64^+^F4/80^+^ cells; monocyte-derived dendritic cell (DC)/macrophage (MΦ) was defined as CD11b^+^CD64^+^Ly6C^+^; Ly6C^+^ monocyte was defined as Ly6C^+^. The cell number of each cell population was calculated using the multiple fluorescent staining image and analyzed by software MetaMorph^TM^, and the data was presented as cell number/ per 664225 (815 × 815). Scale bar is 200 μm. Data are represented as mean ± SEM. \**p*<0.05, \*\**p* < 0.01, \*\*\**p* < 0.001, \*\*\*\**p* < 0.0001 (Student’s *t*-test).

Moreover, severe infiltration of interstitial macrophages (CD11b^+^CD64^+^F4/80^+^), monocyte-derived dendritic cell (DC)/macrophage (MΦ) (CD11b^+^CD64^+^Ly6C^+^), and Ly6C^+^ monocyte (gray column, Fig. 3e) into pulmonary area at day 3 post-SARS-CoV-2 infection were observed in WT mice. In contrast, cell infiltration was attenuated in *clec5a^-/-^tlr2^-/-^* mice (red column, Fig. 3e). These observations indicated that CLEC5A and TLR2 are critical in SARS-CoV-2-induced thromboinflammation in vivo.

### Attenuation of lung collagen deposition in mice deficient of CLEC5A and TLR2

In addition to thromboinflammation, SARS-CoV-2 infection resulted in lung injury and pulmonary fibrosis, including thickening of basement membranes and deposition of collagen [32]. Thus, we were interested to understand whether CLEC5A and TLR2 contributed to SARS-CoV-2-induced collagen deposition. At day 5 post-infection, severe thickening of alveolar cell wall and cell infiltration were noted in WT mice (upper middle, Fig. 4a), while these phenomena were attenuated in *clec5a^-/-^tlr2^-/-^* mice after SARS-CoV-2 infection (upper right, Fig. 4a). We further examined the extent of collagen deposition by Picro Sirius Red Staining. We observed yellow-orange birefringence (type I collagen thick fiber) and green birefringence (type III collagen, thin fiber) in lung tissues of WT mice after SARS-CoV-2 infection (lower middle, Fig. 4a). In contrast, collagen deposition was attenuated in *clec5a^-/-^tkr2^-/-^* mice (lower right, Fig. 4a). These observations suggested that CLEC5A and TLR2 play critical roles in SARS-CoV-2-induced lung fibrosis. The quantiation of collagen area in lung was shown in Fig. 4b. Thus, we concluded that SARS-CoV-2 activated CLEC5A and TLR2 to induce lung inflammatory reactions, thrombosis, and collagen deposition, and PLT-EVs further enhanced NET formation via CLEC5A. These observations further suggested that EVs from virus-activated platelets are potent endogenous danger signals to ehance inflammatory reactions in vivo.

**Figure 4.**
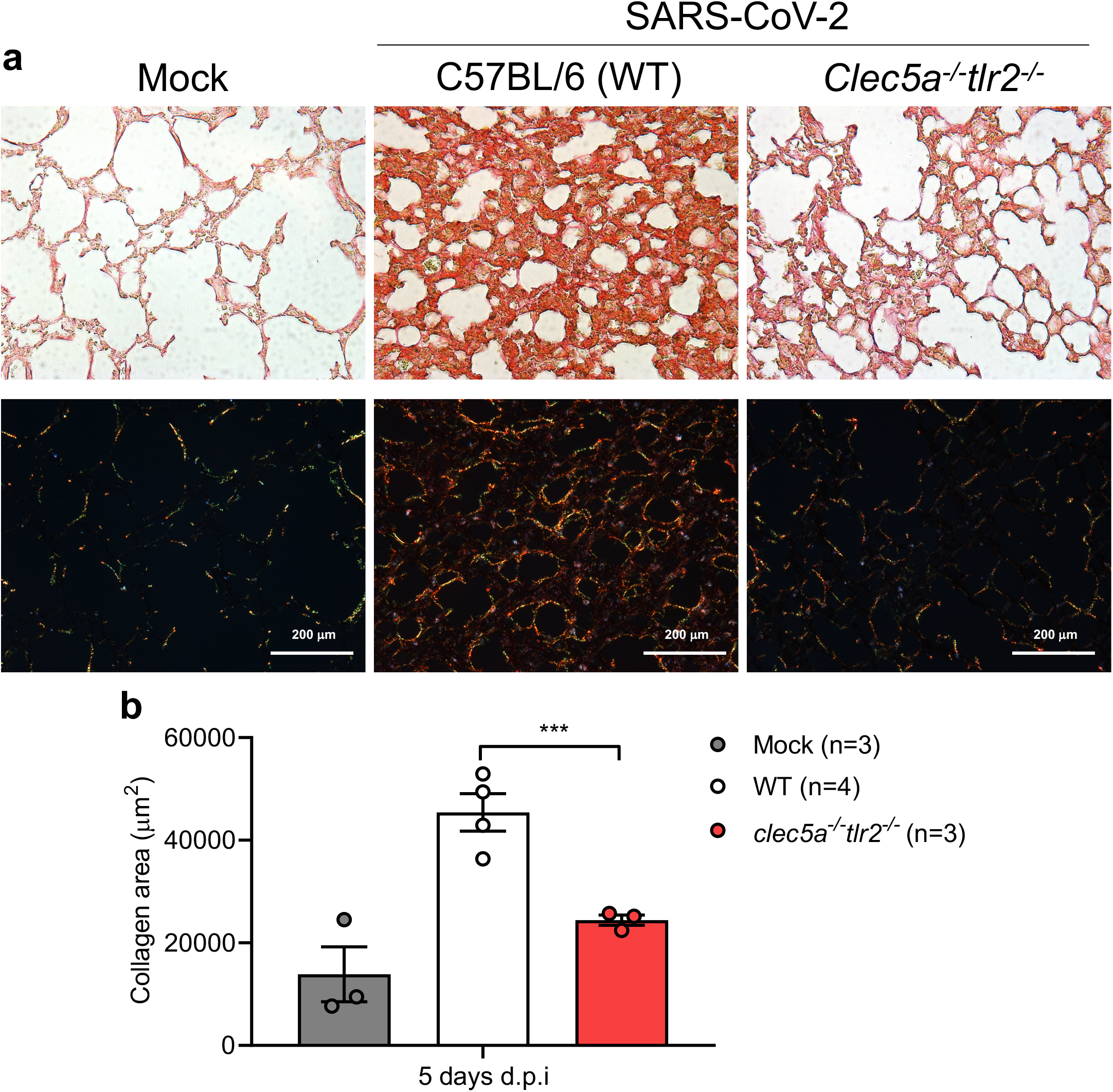
CLEC5A and TLR2 are critical in SARS-CoV-2-induced collagen deposition. SARS-CoV-2-challenged mice were sacrificed to harvest lung at 5 days post-infection. **(a)** Lung sections were stained with Picro Sirius Red dye and captured the image under light microscopy (upper panel) and polarized light microscopy (lower panel) under 25 X magnification. The scale bar is 200 μm. **(b)** The level of collagen deposition was quantified using MetaMorph™ software and presented as area (μm^2^). Data are mean ± SEM. \*\*\**p* < 0.001 (Student’s *t*-test). Data are mean ± SEM. \*\*\**p* < 0.001 (Student’s *t*-test). N=3 of each group.

## Discussion

It has been shown that lung is responsible for 50 % of platelet biogenesis or 10 million platelets per hour [33]. Moreover, platelets were hyperactivated in clinically ill COVID-19 patients [12, 13], and contributed to coagulopathy in COVID-19 patients [17]. Thus, we are interested to understand the mechanism of platelet-mediated immunothrombosis in SARS-CoV-2 infection. In this study, we demonstrated that platelets enhanced SARS-CoV-2-induced NET formation (Supplementary Fig. 1). Moreover, SARS-CoV-2/platelets-induced NETs were detached from the coverslip and floated in the culture medium at 20 h post-incubation, resulting in less NETs attached to coverslip [Supplementary Fig. 2a (lower panel) and 2b (# above white column)]. This phenomenon was not observed in DV/platelets-induced NET formation, suggesting the aggregated floating NETs may contribute to microemboli in COVID-19 patients. Furthermore, SARS-CoV-2 and COVID-19 EVs induced NET formation via CLEC5A and TLR2, and *clec5a^-/-^tlr2^-/-^* mice were resistant to SARS-CoV-2-induced lung inflammation and collagen deposition. These observations suggest that platelets contribute to SARS-CoV-2-induced inflammation significantly, and blockade of CLEC5A and TLR2 may be beneficial to alleviate thromboinflammation and reduce intravascular coagulopathy in COVID-19 patients.

It has been shown that serum EV level correlates with disease severity in COVID-19 patients [34, 35]. Moreover, Syk inhibitor R406 was shown to prevent NETosis of healthy donor neutrophils stimulated with COVID-19 patient plasma [36]. In this study, we further demonstrated that COVID-19 EVs express abundant markers of activated platelets, enhance NET formation via CLEC5A and TLR2 (Figure 3d). These observations are in accord with our previous report that DV activate platelets to release EVs, which are critical endogenous danger signals to trigger NET formation and inflammatory reactions via Syk-coupled CLEC5A and TLR2 [25, 26].

Recently, SARS-CoV-2 was shown to bind ACE2 in platelets to enhance NETosis [17]. However, mass spectrometry [37] and RNA-seq analyses [38, 39] did not detect ACE2 and TMPRSS2 in platelets and megakaryocytes. Increasing evidence indicates that lectins play critical roles in virus-induced systemic inflammation and NET formation [40]. C-type lectins (DC-SIGN and L-SIGN) and sialic acid-binding immunoglobulin-like lectin 1 (SIGLEC1) were shown to function as attachments receptors by enhancing ACE-2 mediated infection [41]. Furthermore, SARS-CoV-2 was reported to exacerbate inflammatory responses in myeloid cells through C-type lectin receptors and Tweety family members [42]. We have shown that DV was captured by DC-SIGN to activate the CLEC2 to release EVs from platelets [25]. Moreover, DV has been shown to be captured by C-type lectins heterocomplex (DC-SIGN and mannose receptor) to trigger inflammation and NETosis via CLEC5A and TLR2 [25, 26, 43]. Thus, it would be very interesting to test whether CLEC2/DC-SIGN complex is responsible for SARS-CoV-2-induced platelet activation in the future.

It has been reported that clinical symptoms, laboratory features and autopsy findings between dengue fever and COVID-19 are similar and are difficult to distinguish, and some patients who were initially diagnosed with dengue, but were later confirmed to have COVID-19 [44]. Moreover, autopsy of COVID-19 patients demonstrated that cells involved in the pathogenesis of COVID-19 and dengue are similar [45]. In this study, we found that platelets and platelet-derived EVs play critical roles in the pathogenesis of SARS-CoV-2. Unlike the thread shape of NET formation caused by DV, SARS-CoV-2-induced robust aggregated NET formation in the presence of platelets. Because the aggregated NETs may detach from SARS-CoV-2-induced thrombi and form micro-emboli in vivo, this observation can explain why intracoagulopathy was only found in COVID-19 patients, but not in DV-infected patients. Thus, inhibition of platelet activation may become a novel strategy to attenuate virus-induced lung inflammatory reactions in the future.

## Supplementary Figure legends

**Supplementary Figure 1.**
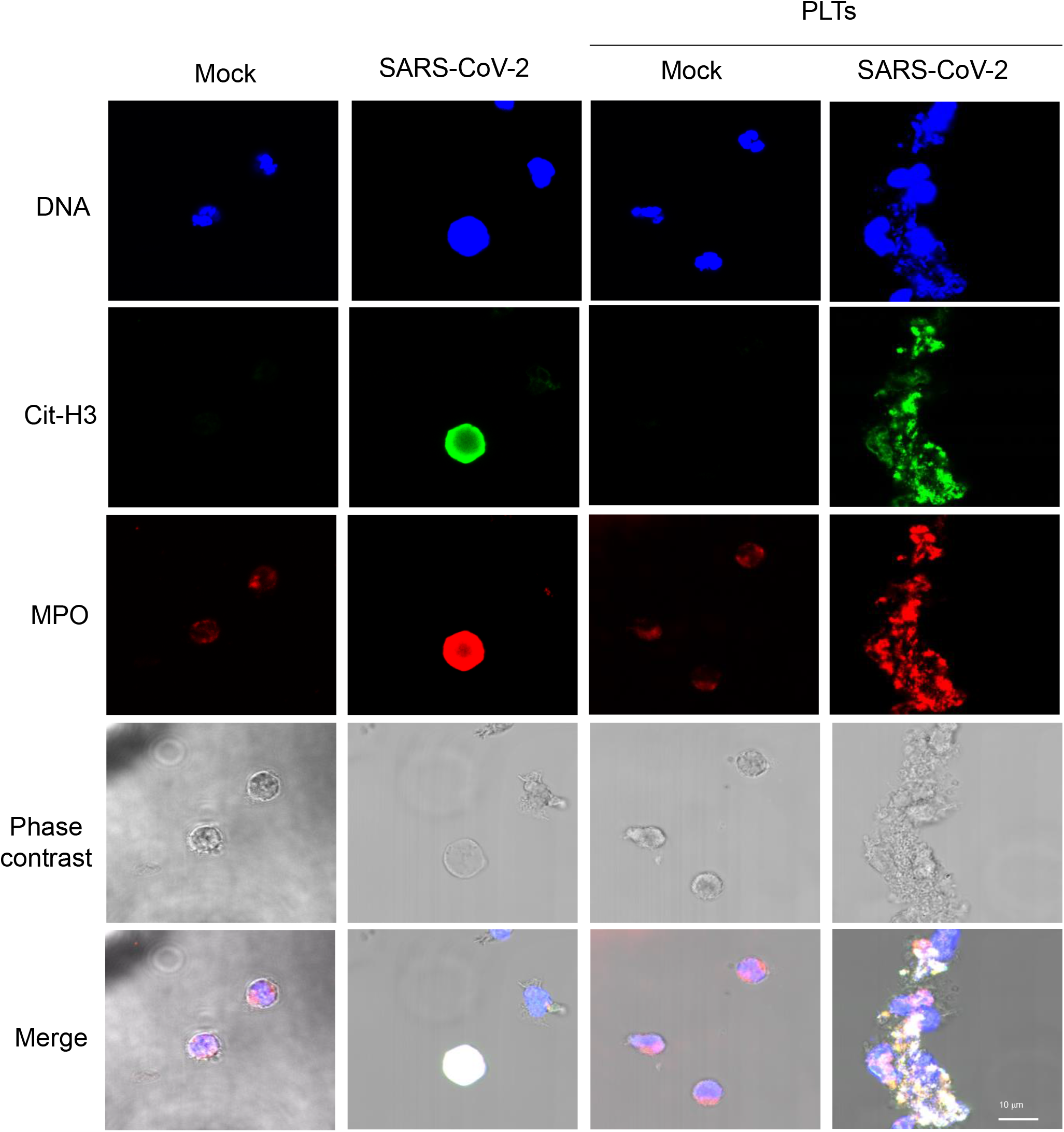
Platelet enhances SARS-CoV-2-induced NET formation. Human neutrophils (4 X10^5^/ml) were incubated with SARS-CoV-2 (MOI=1) in the presence or absence of autologous platelets (4 X10^6^/ml) for 5 h at 37 °C. The NET structure was visualized by staining with DNA (blue), Cit-H3 (green), and MPO (red) then captured by confocal microscopy under 2000 X magnification. The colocalization of DNA, Cit-H3, and MPO was color in white. The scale bar is 10 μm.

**Supplementary Figure 2.**
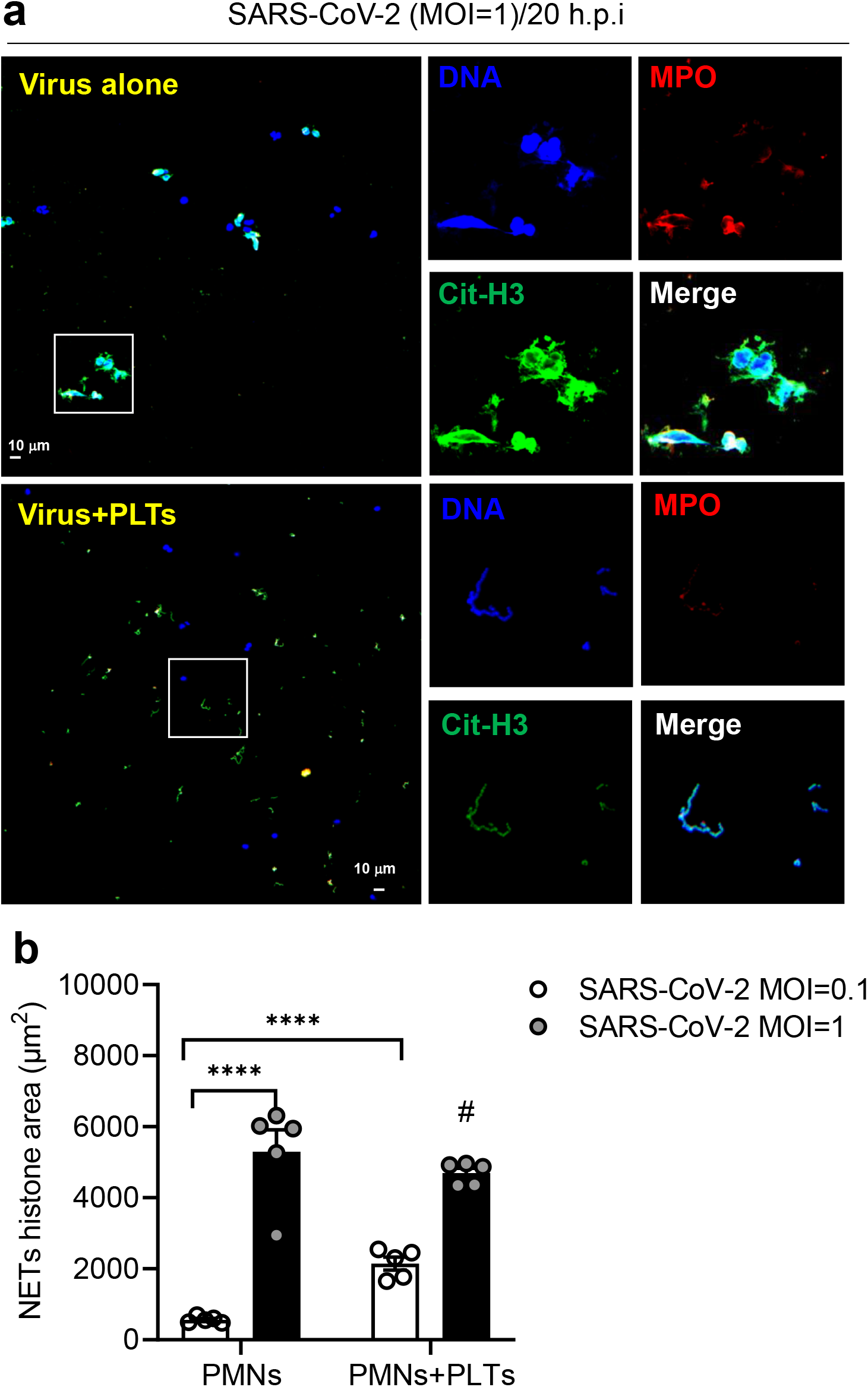
Platelet promotes severe NETs under SARS-CoV-2 stimulation after 20 h incubation. **(a&b)** Neutrophils (4 X10^5^/ml) from healthy volunteers were stimulated with SARS-CoV-2 (MOI=0.1 or 1) with or without autologous platelets (4 X10^6^/ml) for 20 h at 37 ° C. Samples were fixed and stained with DNA (blue), Cit-H3 (green), and MPO (red). The image was captured by confocal microscopy under 400 X magnification (a). The scale bar is 10 μm. The level of NET was calculated using the image of Cit-H3 from 5 healthy donors (b). Data was presented as mean of area (mm^2^) ± SEM. **** *p* < 0.0001 (Student’s *t*-test).

## Acknowledgment

This work was supported by Academia Sinica (AS-IDR-110-01, AS-GC-110-MD01), and Biotechnology Research Park Translational Project (AS-BRPT-110-02). The other supports are from Academia Sinica Investigator Award (AS-IA-109-L02), Ministry of Science and Technology (MOST 107-2321-B-001-015), and VGH, TSGH, AS Joint Research Program (VTA110-V5-5-1). We are grateful to Dr. Yu-Chi Chou to provide SARS-CoV-2 spike pseudotyped virus, and Dr. Jia-Tsrong Jan for technical support.

## Human samples

This study was conducted under the Helsinki Declaration of 1975. All patients provided written informed consent before enrollment, and the study was approved by the Research Ethics Committee of Academia Sinica, Taipei (AS-IRB01-20024, AS-IRB01-20025).

## Statement of animal operation in BSL-3 biosafety animal facility

All mouse works were conducted in accordance with the “Guideline for the Care and Use of Laboratory Animals” as defined by the Council of Agriculture, Taiwan. Mouse work was approved by the Institutional Animal Care and Use Committee of Academia Sinica (protocol ID: 20-05-1471 and 19-07-1330). The Institutional Biosafety Committee of Academia Sinica approved work with infectious SARS-CoV-2 virus strains under BSL3 conditions. All sample processes were conducted according to “Interim Laboratory Biosafety Guidelines for Handling and Processing Specimens Associated with Coronavirus Disease 2019 (COVID-19)” recommended by CDC.

